# Comparison between three concentration techniques for diagnosing intestinal parasites

**DOI:** 10.1101/308015

**Authors:** Sonia Trabelsi, Majdi Hanachi, Sarra Cheikhrouhou, Dorsaf Aloui, Meriam Bouchekoua, Samira Khaled

**Affiliations:** Parasitology and Mycology Department, Charles Nicolle University Hospital, Tunis, Tunisia

**Keywords:** Parasitic intestinal diseases, Diagnostic techniques and procedures, Sensitivity

## Abstract

**Background:** Intestinal parasitoses still are a noticeable threat to public health. The direct diagnosis of such parasites requires the use of concentration techniques, whose sensitivities for protozoan cysts and helminth eggs are far from equal.

**Aim:** To compare the Willis, Ritchie and Bailenger concentration techniques in terms of parasite recovery, cost, time, and biosafety.

**Methods:** This prospective study analysed 236 stool specimens for intestinal parasites using the direct wet smear and the above-mentioned concentration techniques applied separately.

**Results:** Biphasic techniques identified significantly more positive specimens for intestinal parasites than the Willis technique, the latter leading to less concentrated and more altered parasitic elements on microscopy. No statistically significant difference emerged from comparing Ritchie’s and Bailenger’s methods. The Willis technique was the safest, yet the costliest and the most time-consuming of the studied methods.

**Conclusions:** Even though the hazardous reagents employed may raise legitimate concerns over their health implications, biphasic techniques prove to be uncostly, quick to perform, and highly sensitive for detecting faecal parasites, therefore ensuring a safe diagnosis for routine stool examinations.

## INTRODUCTION

Human intestines and biliary ducts can host a wide range of saprophytic and parasitic organisms. Some of the latter may turn out to be pathogenic, causing intestinal parasitoses. The main mode of transmission of such diseases is the faecal-oral route. Despite the significant improvement in terms of hygienic conditions and the subsequent decrease in their incidence, these pathologies should not be relegated to the background. In fact, they still constitute a major public health problem in many developing countries, leading to a noticeable morbimortality and a negative impact on their economy (1).

Since symptoms are not specific, the diagnosis of intestinal parasitoses cannot be established clinically and needs to be confirmed by further tests. In this regard, the laboratory plays a crucial role in diagnosing parasitic intestinal infections, mostly through a parasitological stool examination. This test must include a direct wet smear and a direct microscopic examination after performing a concentration technique (2), as decreed by the Tunisian ministry of public health in the nomenclature of clinical pathology acts.

Several stool concentration methods were developed throughout the years, applying different chemical and physical principles. Biphasic techniques, combining the action of chemical reagents with a physical process, appear to be the most widely used nowadays, especially resource-poor countries (3). No technique can guarantee the recovery of all parasites present in a faecal sample, each method being characterized by its advantages and its limits. To deal with this issue in the absence of standardisation, some laboratories resort to the use of two complementary methods in order to optimize their results. Other criteria are to be taken into account when evaluating a concentration technique, such as its cost and the toxicity of the reagents it employs. These criteria are critical in the context of developing countries, which happen to be the most affected by intestinal parasitoses, as concentration techniques must ally efficiency and affordability without violating the biosafety standards.

The aim of the present study was to compare between three parasite concentration techniques, namely the Willis, Ritchie and Bailenger methods, based on sensitivity, time of realisation, cost, and biosafety.

## MATERIALS AND METHODS

### Stool examination procedure

This prospective study encompassed 236 faecal specimens coming from outpatients, inpatients, or non-permanent resident students in Tunisia.

A direct wet smear was performed by spreading a small amount of the sample with a drop of physiological serum before applying a coverslip. The whole smear thus obtained was examined with the low-power objective (10x), while the high-power objective (40x) was used to observe selected fields (2). The evaluated concentration techniques were then performed on separate samples of each faecal specimen as detailed below.

#### Willis concentration technique

Two grams of stool were diluted in 20 milliltres of sodium chlorate. The dilution was homogenised. The solution was strained through. The obtained suspension was poured into a tube until its superior limit (a mild bombing of the liquid above the border). A coverslip was then delicately applied on top of the tube while avoiding air bubbles. A quarter of an hour later, the coverslip was removed and deposited on a microscope slide (4).

#### Ritchie concentration technique

The stool sample was diluted in 10% formalin in water. The mixture was strained through two layers of gauze into a conical 30-ml centrifuge tube until 30 ml. The tube was centrifuged for 2 minutes at 1500 revolutions per minute (rpm). The supernatant fluid was decanted and discarded. The remaining faecal sediment was thoroughly mixed with 10% formalin. The tube was filled with 10% formalin until 20 ml, then with diethyl ether until 30 ml. The tube was stoppered and vigorously shaken for homogenisation. A second centrifugation was performed with the same parameters. Four layers were obtained: a top layer of ether, a debris plug layer, a formal saline layer and a sediment layer in the bottom of the tube. The upper layers were eliminated by quickly inverting the tube. Two drops of the remaining sediment were deposited on a microscope slide with a Pasteur pipette. A coverslip was added before examining the slide with a microscope (5).

#### Bailenger concentration technique

Two point five grams of the stool sample were diluted in 25 ml of aceto-acetic buffer. The mixture was then sieved using a gauze and collected in a conical tube until reaching a volume of 20 ml. The same volume of ether is added before strenuously shaking the mixture. After centrifuging the tube for one minute at 1500 rpm and decanting the supernatant, a few drops of the sediment were deposited on a microscope slide and overlaid with a coverslip for microscopic examination (6).

### Statistical analysis

We compared the concentration techniques’ performances in identifying faecal parasites using McNemar test for paired samples. The Statistical Package for the Social Sciences (SPSS) version 22.0 software was used to calculate all the parameters. Differences were considered statistically significant if *P* values were < 0.05 and highly significant if *P* values were < 0.001.

## RESULTS

Of the 236 faecal specimens included in our study, intestinal parasites were detected by the direct wet smear and/or the microscopic examination after concentration in 79 samples, which means a global prevalence of 33.47%. Parasites were detected by the direct wet smear in 88.6% of the positive specimens, while only 57% of the latter were identified thanks to the concentration methods employed (table 1). *Blastocystis sp*. and *Dientamoeba fragilis* were detected almost only using the direct wet smear. Protozoan cysts and helminth ova were mainly identified after performing a concentration technique (table 2).

**TABLE 1.** Positive samples according to the direct wet smear versus concentration techniques (all techniques included)

|  |  | Direct wet smear |  | Total |
| --- | --- | --- | --- | --- |
|  |  | Positive | Negative |  |
| Concentration techniques | Positive | 36 | 9 | 45 |
|  | Negative | 34 | 157 | 191 |
| Total |  | 70 | 166 | 236 |

**TABLE 2.**
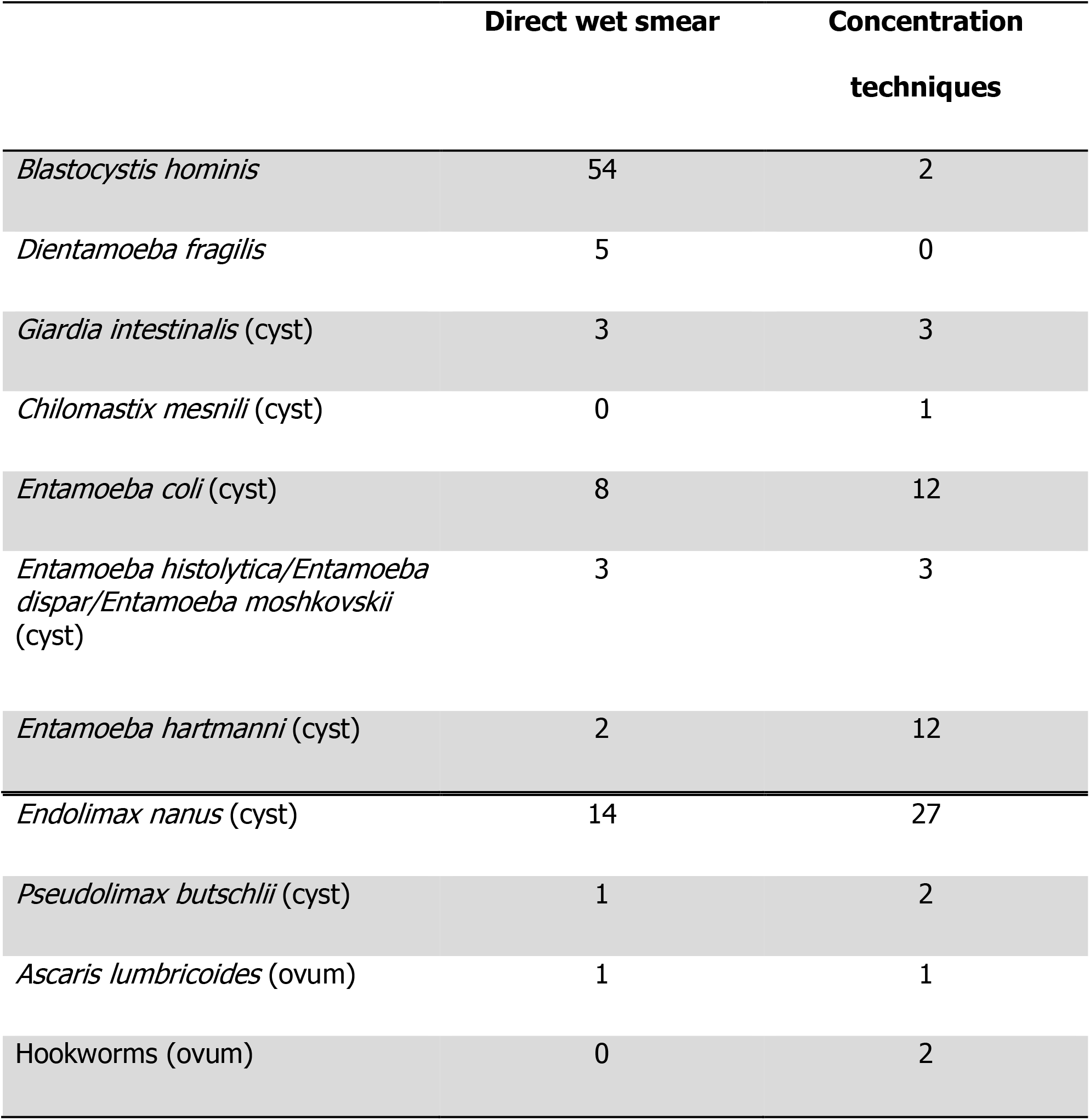
Intestinal parasites identified by the direct wet smear versus recovered by concentration techniques (all techniques included)

### Comparison of parasite recovery

Intestinal parasites were detected in 11 samples (14.1% of positive specimens) using the Willis flotation technique, while Bailenger’s and Ritchie’s biphasic methods were able to identify protozoan cysts and/or helminth ova in 42 (53.84%) and 44 (56.41%) samples respectively. Table 3 compares the Willis technique to the biphasic ones. The latter are in turn compared in table 4. The number of positive samples per parasitic species according to each concentration technique is presented in table 5.

**TABLE 3.**
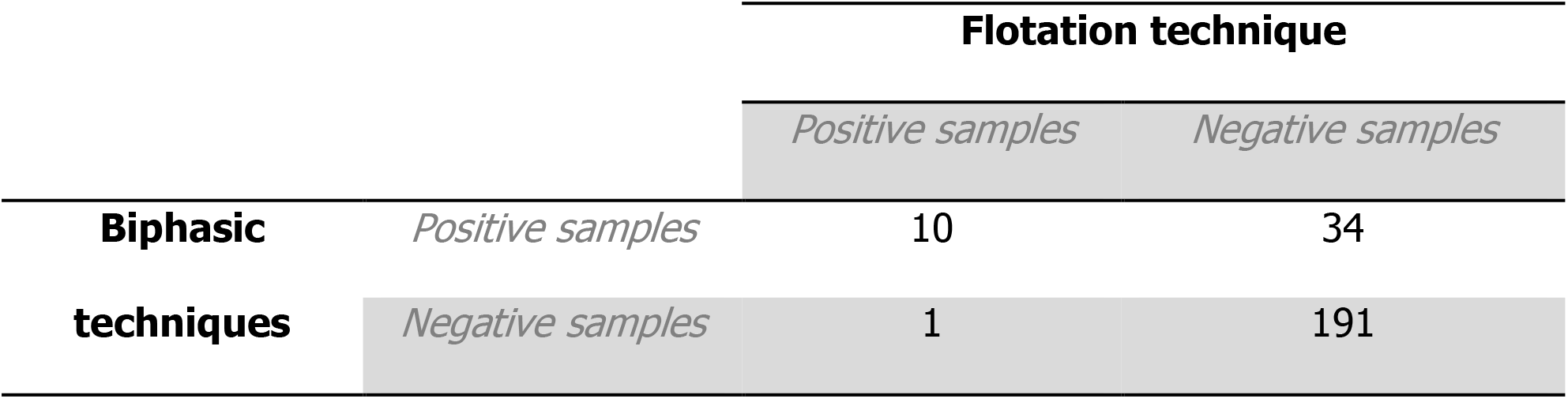
Comparison between a flotation technique (Willis’) and two biphasic techniques (Ritchie’s and Bailenger’s) for the diagnosis of intestinal parasites in human faecal samples

**TABLE 4.**
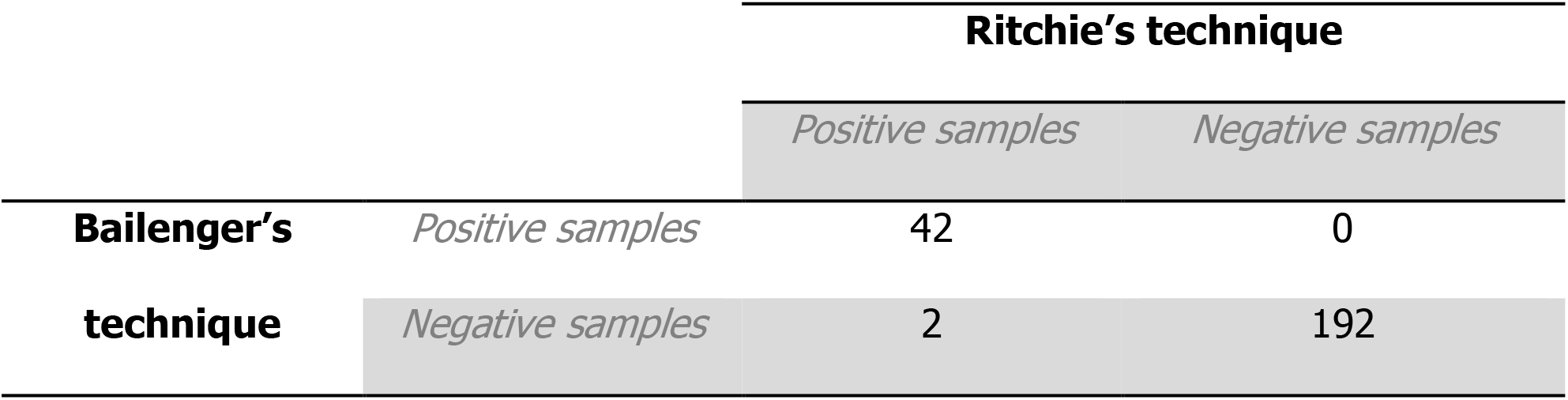
Comparison between Ritchie’s and Bailenger’s concentration techniques for the diagnosis of intestinal parasites in human faecal samples

**TABLE 5.** Number of positive human stool specimens per intestinal parasite according to three different concentration techniques

| Parasitic species | Willis | Ritchie | Bailenger |
| --- | --- | --- | --- |
| *Entamoeba coli* (cyst) | 2 | 12 | 12 |
| *Entamoeba histolytica*/<br>*Entamoeba dispar*/<br>*Entamoeba moshkovskii*<br>(cyst) | 0 | 3 | 3 |
| *Entamoeba hartmanni* (cyst) | 2 | 12 | 12 |
| *Endolimax nanus* (cyst) | 3 | 27 | 25 |
| *Pseudolimax butschlii* (cyst) | 1 | 2 | 2 |
| *Giardia intestinalis* (cyst) | 2 | 3 | 3 |
| *Chilomastix mesnili* (cyst) | 0 | 1 | 1 |
| *Ascaris lumbricoides* (ovum) | 1 | 1 | 1 |
| Hookworms (ovum) | 2 | 2 | 2 |

By applying the McNemar test for paired samples, the following results were obtained: Willis versus biphasic techniques (*P* < 0,001): the difference is statistically highly significant; Bailenger versus Ritchie (*P* = 0.5): the difference is not statistically significant.

Other microscopic parameters were analysed, such as the abundance of parasitic elements on microscopy (table 6) as well as the degree of conservation of their morphology (table 7). It thus appears that the flotation technique not only fails to recover as much parasites as the biphasic techniques, but also alters the eggs’ shell. In our study, Ritchie’s method recovered more protozoan cysts than Bailenger’s.

**TABLE 6.** Comparison of the abundance of parasites recovered between three concentration techniques

|  | Willis | Bailenger | Ritchie |
| --- | --- | --- | --- |
| **Cysts** | + | ++ | +++ |
| **Eggs** | + | +++ | +++ |

+: a few parasitic elements; ++: moderately rich; +++: very rich

**TABLE 7.** Comparison of the conservation of parasites recovered between three concentration techniques

|  | **Willis** | **Bailenger** | **Ritchie** |
| --- | --- | --- | --- |
| **Cysts** | — | + | + |
| **Eggs** | — | + | + |

—: alteration; +: integral conservation

### Comparison of cost

Table 8 exposes the cost of each technique by calculating the price in the Tunisian market as of November 2017 of all the material needed to concentrate the 236 samples included in the study. While Ritchie’s method is the cheapest, closely tailed by Willis’ flotation procedure, the Bailenger technique’s expensiveness can be explained by the use of larger measuring tubes and a greater quantity of ether.

**TABLE 8.** Cost in US dollars of reagents and material required by each of the three studied techniques to concentrate 236 specimens (Tunisian market prices in November 2017)

|  | **Willis** | **Bailenger** | **Ritchie** |
| --- | --- | --- | --- |
| Sodium chloride | 3.513 | - | - |
| Crystallized sodium acetate | - | 1.573 | - |
| Acetic acid | - | 0.172 | - |
| Formaldehyde | - | - | 1.660 |
| Ether | - | 21.365 | 10.682 |
| Gloves | 15.708 | 15.708 | 15.708 |
| Pasteur pipettes | - | 6.813 | 6.813 |
| Microscope slides | 1.514 | 1.514 | 1.514 |
| 24- by 24-mm coverslip | 2.366 | 2.366 | 2.366 |
| Conical tubes | 12.112 | - | 12.112 |
| Measuring tubes (50 ml) | 28.104 | 56.209 | 28.104 |
| Measuring tubes (15 ml) | 19.399 | - | - |
| Graduated pipettes | - | 0.734 | - |
| pH paper | - | 0.191 | - |
| Wooden sticks | 0.237 | 0.237 | 0.473 |
| **Total** | 82.953 | 106.882 | 79.432 |

### Comparison of time

As shown by table 9, which compares the required time to perform each of the studied techniques, Bailenger’s method is the fastest to perform, while the Willis concentration technique requires more than twice as much time than the former technique.

**TABLE 9.** Mean time needed *per* parasite concentration method

|  | **Willis** | **Bailenger** | **Ritchie** |
| --- | --- | --- | --- |
| **Mean time** | 18 minutes | 8 minutes and 5 seconds | 13 minutes and 4 seconds |

## DISCUSSION

The aim of this study was to compare three concentration techniques not only in terms of parasite recovery, but also according to practical criteria such as cost, processing time, and biosafety. We thus evaluated a flotation method, the Willis technique, and two biphasic methods, the Ritchie and the Bailenger techniques. This choice was motivated by the fact that these concentration methods are the most frequently used in parasitology laboratories in developing countries (3).

As for parasite recovery, concentration techniques were more efficient than the direct wet smear for identifying protozoan cysts and helminth ova, this performance being the reason why the use of these techniques is mandatory in routine stool examinations. On the other side, we mainly rely on the direct wet smear for diagnosing *Blastocystis sp*. and *Dientamoeba fragilis*, too fragile to be observed after performing a concentration method. Our results match those obtained by Oguoma *et al*. (3), the prevalence of helminth and protozoa detected by a formol-ether concentration technique being significantly higher than the one found by the direct smear.

The comparison of sensitivity between the concentration techniques included in this study showed a statistically highly significant difference in favour of the biphasic methods. Even though Ritchie’s method recovered slightly more parasites than Bailenger’s, the difference was not statistically significant. On the microscopic level, the comparative analysis did also highlight more abundant and better conserved parasites when using the biphasic techniques compared to the flotation method. The latter may therefore fail to diagnose intestinal parasites because of their limited number in the sample or due to an altered morphology that would render them unrecognisable. In agreement with our work, Bartlett *et al*. (7) drew the conclusion that the formalin-ether concentration method was more efficient than the modified zinc sulfate flotation technique it was compared to.

This study did also demonstrate a distinctive superiority of biphasic techniques over Willis’ method for routine stool examination, since the latter proved to be more expensive and more time-consuming. These criteria, along with sensitivity, are important to take into consideration when picking the concentration technique to perform on a daily basis in the laboratory. On a larger scale, the need for simple and cheap yet efficient concentration techniques is crucial in developing countries — which also happen to be endemic for numerous intestinal parasites — in order to adapt to the cost containment policies in public health.

However, Bailenger’s and Ritchie’s techniques resort to hazardous reagents in their procedure. In fact, ether, employed by both above-mentioned methods, is flammable and irritating to skin, eyes and upper respiratory system (8). Symptoms induced by acetic acid, used as a fixative by the Bailenger method, vary from conjunctivitis and throat irritation to skin and eye burns (9). Ritchie’s technique relies on formalin, another irritating reagent and a potential carcinogen after chronic exposure (10). No reagent employed by the Willis method has any kind of chemical hazards that may endanger the laboratory staff. Some authors demonstrated that less toxic reagents could be used in replacement of ether as a solvent to extract fat and debris, like ethyl acetate (11), acetone (12), or tween (13). A modified version of Ritchie’s method by Régis Anécimo (14) did even replace both formaldehyde and ether by a natural detergent, yet had similar qualitative and quantitative performances in parasite recovery. Some protocols resort to sodium hydroxide (NaOH) to perform the formol-ether concentration technique, but a comparative study conducted by Suwansaksri et al. (15) found no statistically significant difference when comparing its detection rate with a normal saline preparation, allowing to avoid the use of NaOH for security reasons. Laboratory technicians should therefore be aware of the health implications the use of biphasic techniques exposes to in order to strictly comply with the appropriate biosafety measures.

## CONCLUSIONS

Biphasic techniques proved their superiority over Willis’ flotation technique as they happen to be uncostly, quick to perform, and highly sensitive for detecting intestinal parasites, whether it be protozoan cysts or helminth ova. Even though the hazardous reagents employed may raise legitimate concerns over their health implications, these techniques ensure a reliable diagnosis for routine laboratory analysis.

In the absence of any international or national recommendation, conducting comparative studies between concentration techniques would be interesting for any laboratory in order to evaluate the affordable methods based on objective criteria, leading to the implementation of the fittest technique in the daily routine protocols. Charles Nicolle Teaching Hospital’s parasitology and mycology laboratory proceeded this way before picking Ritchie’s method, whose qualities were demonstrated by the present study, among others (16).

In the light of health issues that such techniques give rise to, further inquiries should look for safer intestinal parasite concentrators that would be at least as efficient. Evaluating commercial kits in comparison to in-home biphasic techniques would be valuable, particularly since the promotion of these alternatives focuses on biosafety guarantees.

## Acknowledgement

This work forms part of the end of studies project of R. Guenichi and R. Hanchi, under the supervision of S. Trabelsi, MD, Professor of Medicine.

This research received no specific grant from any funding agency in the public, commercial, or not-for-profit sectors. The authors declare that they do not have any commercial associations which might give rise to a conflict of interest in connection with the submitted article.

## REFERENCES

1. Haque R. 2007. Human Intestinal Parasites. J Health Popul Nutr 25(4):387–91.

2. Garcia L. 2007. Macroscopic and Microscopic Examination of Fecal Specimens, p 782–830. *In* Garcia L (ed), Diagnostic Medical Parasitology, 5th ed. ASM Press, Washington, DC.

3. Oguoma VM, Ekwunife CA. 2007. The Need For A Better Method: Comparison Of Direct Smear And Formol-Ether Concentration Techniques In Diagnosing Intestinal Parasites. Internet J Trop Med, vol 3, number 2.

4. Willis HH. 1921. A Simple Levitation Method for the Detection of Hookworm Ova. Med J Aust 2:375–76.

5. Ritchie LS. 1948. An ether sedimentation technique for routine stool examinations. Bull US Army Med Dep 8(4):326.

6. Bailanger J. 1965. Parasitological and functional coprology (in French). Imprimerie E. Drouillard, Bordeaux.

7. Bartlett MS, Harper K, Smith N, Verbanac P, Smith JW. 1978. Comparative evaluation of a modified zinc sulfate flotation technique. J Clin Microbiol 7(6):524–8.

8. National Institute for Occupational Safety and Health. NIOSH Pocket Guide to Chemical Hazards. Third printing. 2007. Ethyl ether; p. 140.

9. National Institute for Occupational Safety and Health. NIOSH Pocket Guide to Chemical Hazards. Third printing. 2007. Acetic acid; p. 2.

10. National Institute for Occupational Safety and Health. NIOSH Pocket Guide to Chemical Hazards. Third printing. 2007. Formaldehyde; p. 148.

11. Truant AL, Elliott SH, Kelly MT, Smith JH. 1981. Comparison of formalin-ethyl ether sedimentation, formalin-ethyl acetate sedimentation, and zinc sulfate flotation techniques for detection of intestinal parasites. J Clin Microbiol 13(5):882–84.

12. Feleke M, Yeshambel B, Moges T, Yenew K, Andargachew M, Afework K, Kahsay H. 2010. Comparison of formol-acetone concentration method with that of the direct iodine preparation and formol-ether concentration methods for examination of stool parasites. Ethiop J Health Dev 24(2):148–51.

13. Methanitikorn R, Sukontason K, Sukontason KL, Piangjai S. 2003. Evaluation of the formalin-Tween concentration technique for parasitic detection. Rev Inst Med Trop Sao Paulo 45(5):289–91

14. Anécimo RS, Tonani KAA, Fregonesi BM, Mariano AP, Ferrassino MDB, Trevilato TMB, Rodrigues RB, Segura-Muñoz SI. 2012. Adaptation of Ritchie’s method for parasites diagnosing with minimization of chemical products. Interdiscip Perspect Infect Dis 2012:409757.

15. Suwansaksri J, Nithiuthai S, Wiwanitkit V, Soogarun S, Palatho P. 2002. The formol-ether concentration technique for intestinal parasites: comparing 0.1 N sodium hydroxide with normal saline preparations. Southeast Asian J Trop Med Public Health 33 Suppl 3:97–8.

16. Sato C, Rai SK, Uga S. 2014. Re-evaluation of the formalin-ether sedimentation method for the improvement of parasite egg recovery efficiency. Nepal Med Coll J 16(1):20–5

